# Neuromusculoskeletal simulation reveals abnormal rectus femoris-gluteus medius reflex coupling in post-stroke gait

**DOI:** 10.1101/501072

**Authors:** Tunc Akbas, Richard R. Neptune, James Sulzer

**Author notes:** **Correspondence**: James Sulzer.

## Abstract

Post-stroke gait is often accompanied by muscle impairments that result in adaptations such as hip circumduction to compensate for lack of knee flexion. Our previous work robotically enhanced knee flexion in individuals post-stroke with Stiff-Knee Gait (SKG), however, this resulted in greater circumduction, suggesting the existence of abnormal coordination in SKG. The purpose of this work is to investigate two possible mechanisms of the abnormal coordination: 1) an involuntary coupling between stretched quadriceps and abductors, and 2) a coupling between volitionally activated knee flexors and abductors. We used previously collected kinematic, kinetic and EMG measures from nine participants with chronic stroke and five healthy controls during walking with and without the applied knee flexion torque perturbations in the pre-swing phase of gait in the neuromusculoskeletal simulation. The measured muscle activity was supplemented by simulated muscle activations to estimate the muscle states of the quadriceps, hamstrings and hip abductors. We used linear mixed models to investigate two hypotheses: H1) association between quadriceps and abductor activation during an involuntary period (reflex latency) following the perturbation and H2) association between hamstrings and abductor activation after the perturbation was removed. We observed significantly higher rectus femoris (RF) activation in stroke participants compared to healthy controls within the reflex latency period following the perturbation based on both measured (H1, p < 0.001) and simulated (H1, p = 0.022) activity. Simulated RF and gluteus medius (GMed) activations were correlated only in those with SKG, which was significantly higher compared to healthy controls (H1, p = 0.030). There was no evidence of voluntary synergistic coupling between any combination of hamstrings and hip abductors (H2, p > 0.05) when the perturbation was removed. The RF-GMed coupling suggests an underlying abnormal reflex coordination pattern in post-stroke SKG. These results challenge earlier assumptions that hip circumduction in stroke is simply a kinematic adaptation due to reduced toe clearance. Instead, abnormal coordination may underlie circumduction, illustrating the deleterious role of abnormal coordination in post-stroke gait.

## 1 INTRODUCTION

Following a stroke, individuals often experience significant impairments including muscle weakness, spasticity, increased tone, and abnormal coordination (Twitchell 1951), which often results in compensatory movements(Perry and Burnfield 1992). Abnormal coordination has been quantified in the upper limb (Dewald, Pope et al. 1995),but only more recently in the lower limbs using joint torque measures (Cruz and Dhaher 2008, Neckel, Blonien et al. 2008, Cruz, Lewek et al. 2009, Sakuma, Ohata et al. 2014), mechanical perturbations (Finley, Perreault et al. 2008, Sakuma, Ohata et al. 2014) and H-reflex stimulations (Marque, Simonetta-Moreau et al. 2001, Maupas, Marque et al. 2004, Dyer, Maupas et al. 2009, Dyer, Maupas et al. 2011). However, it is unclear whether such abnormal coordination has direct consequences on gait function. Descriptive gait analyses based on biomechanical data and simulated muscle activations during gait in post-stroke individuals (Hidler, Carroll et al. 2007, Clark, Ting et al. 2009, Lauziere, Betschart et al. 2014) and gait in cerebral palsy (Steele, Rozumalski et al. 2015, Shuman, Schwartz et al. 2017) suggest that lack of lower limb coordination correlates with gait dysfunction. Yet descriptive analysis lacks the ability to identify the causal mechanisms of abnormal coordination that perturbation methods can provide.

Our previous work developed a device that delivers controlled knee flexion torque perturbations during gait (Sulzer, Roiz et al. 2009) and applied it to individuals post-stroke with Stiff-Knee Gait (SKG) (Sulzer, Gordon et al. 2010). SKG is defined by reduced knee flexion during the swing phase, often assumed to be compensated for with hip circumduction (Perry and Burnfield 1992). However, when exposed to pre-swing knee flexion torque perturbations, those with SKG walked with exaggerated hip abduction during swing instead of the expected reduction, while there was no change in healthy controls (Sulzer, Gordon et al. 2010). Biomechanical factors such as balance and perturbation dynamics could not account for the increased abduction. Thus, abnormal neural behavior appears to underlie the hip abduction in people with SKG. However, the neural mechanisms leading to the exaggerated hip abduction remain unclear. The abnormal cross-planar kinematics were likely the result of abnormal heteronymous muscular activation initiated via reflexive (Finley, Perreault et al. 2008) or voluntary (Cruz and Dhaher 2008, Tan and Dhaher 2014) mechanisms. For instance, cross-planar reflexive couplings between adductor longus (AL) and RF were observed in individuals post-stroke while in a seated position (Finley, Perreault et al. 2008). Also in post-stroke individuals, voluntary activation of the hamstrings resulted in a cross-planar coupling with adductors while standing (Cruz and Dhaher 2008). This is in agreement with previous simulation analyses of post-stroke gait showing synergetic coupling between hip abductors and knee flexors in the swing phase (Allen, Kautz et al. 2013). Based on this information, the exaggerated abduction observed in our previous work may have been the result of a reflexive stimulus of the RF coupled with abductor muscles (Hypothesis 1). In contrast, such abductor activity may have been coupled with voluntary initiation of the hamstrings (Hypothesis 2) as an adaptation to the perturbations.

In this work, we seek to delineate the underlying muscle activation behind the abnormal cross-planar kinematics. We used neuromusculoskeletal modeling and simulation (NMMS) to derive the estimated muscle states and supplement measured EMG. NMMS combines the use of a lower limb musculoskeletal model with measured kinematic and kinetic data to estimate muscle states (Zajac 2002, Zajac, Neptune et al. 2003). Descriptive NMMS simulations have identified the role of rectus femoris in post-stroke gait (Reinbolt, Fox et al. 2008). Here we used NMMS to estimate muscle states that are difficult to measure experimentally during gait, such as fiber stretch velocity and all lower limb muscle activities, including abductor muscles that are difficult to access using EMG. Thus, together with measured EMG, NMMS can help elucidate mechanisms of cross-planar muscle synergies and abnormal reflexive responses. Evidence of an abnormal reflex coupling underlying excessive hip abduction during knee flexion perturbations in those with SKG would manifest itself as a hyperactive RF stretch reflex followed by heteronymous activation in the hip abductors (H1). Alternatively, if the abnormal coupling was generated by the voluntary knee flexion movement and due to lack of independent joint control, temporary removal of torque perturbations (“catch trials”) should result in correlated activation between the hamstrings and abductors (H2). This study represents a novel approach towards delineating the differential roles of impairments in post-stroke gait.

## 2 METHODS

### 2.1 Experimental Data

Nine chronic, hemiparetic participants with post-stroke SKG and five healthy, unimpaired individuals gave written informed consent using procedures approved by the local Institutional Review Board to participate in the experiment and described in detail in previous work(Sulzer, Gordon et al. 2010). Inclusion criteria for the hemiparetic participants included reduced knee flexion angle during swing and the ability to walk for 20 minutes without rest at 0.55 m/s on a treadmill (Sulzer, Gordon et al. 2010). A lightweight, custom-designed powered knee orthosis was used to provide knee flexion torque perturbations during the pre-swing phase without affecting the remainder of the gait cycle (Sulzer, Roiz et al. 2009). The level of torque perturbations was automatically adjusted to maximize the swing phase knee flexion angle in those with SKG, whereas in healthy controls the torque was adjusted to increase knee flexion angle at 20° during the swing for each subject prior to commencing the experiment.

The protocol consisted of 610 steps and lasted 16 minutes. No perturbation was applied during the initial 50 steps (baseline) and the perturbation was applied for the next 560 steps (perturbed). Also, four non-consecutive trials (catch trials) with no perturbation during a single gait cycle were implemented throughout the perturbation period. Lower limb kinematic data were collected using optical motion capture (Motion Analysis, Santa Rosa, CA), ground reaction forces were collected using an instrumented treadmill (Tecmachine, Andrez Boutheon, France), and the applied perturbation torque was collected through the knee orthosis. Surface EMG (Delsys Inc., Boston, MA) data were collected from RF, vastus lateralis, adductor longus, medial hamstrings in healthy participants and additionally the tensor fascie latae (TFL), and vastus medialis in hemiparetic patients. Motion capture data were collected at 120 Hz, whereas other data were collected at 1 kHz.

### 2.1 EMG Analysis

EMG data were used to determine reflex activation and validate the NMMS model. All signal processing of EMG data was performed in MATLAB (MathWorks, Natick, MA). To evaluate reflex activation, each EMG signal was demeaned, rectified and then integrated over a temporal window 100 ms prior to toe-off, which was the window expected to reveal stretch reflex response to perturbations (Mrachacz-Kersting, Lavoie et al. 2004). We analyzed 10 and 20 consecutive steps from baseline and perturbation periods, respectively. For NMMS validation, the EMG signals were demeaned, rectified and low-pass filtered with a 5th order Butterworth filter at 20 Hz. The processed signals were then synced with kinematic data, divided into gait cycles and integrated into 1% gait cycle intervals to obtain integrated EMG (iEMG) measures. The magnitude of iEMG was normalized to the participant’s own maximum overall iEMG across all trials.

### 2.2 Neuromusculoskeletal Model and Simulation

We used NMMS to determine muscle states from kinematic, kinetic, and perturbation data with the Gait 2392 musculoskeletal model of OpenSim version 3.3 (Delp, Anderson et al. 2007). The model consisted of 18 degrees-of-freedom and 80 muscles. We condensed the head, arms and trunk (HAT) to the pelvis segment in the model since no markers were placed on the HAT. For each participant, the joint angles were calculated using inverse kinematics with the least square fit of marker trajectories. The residual reduction algorithm (RRA) was used to fine-tune the model to minimize the residuals (artificial forces and moments to maintain dynamic consistency of the simulation) between the kinematic and GRF data. The parameters required to test our hypotheses, i.e. muscle activation profiles and muscle fiber stretch values, were calculated using computed muscle control (CMC). CMC identifies the muscle activation patterns to produce a forward dynamics simulation emulating the experimentally collected data. Figure 1 illustrates the NMMS procedure.

**Figure 1.**
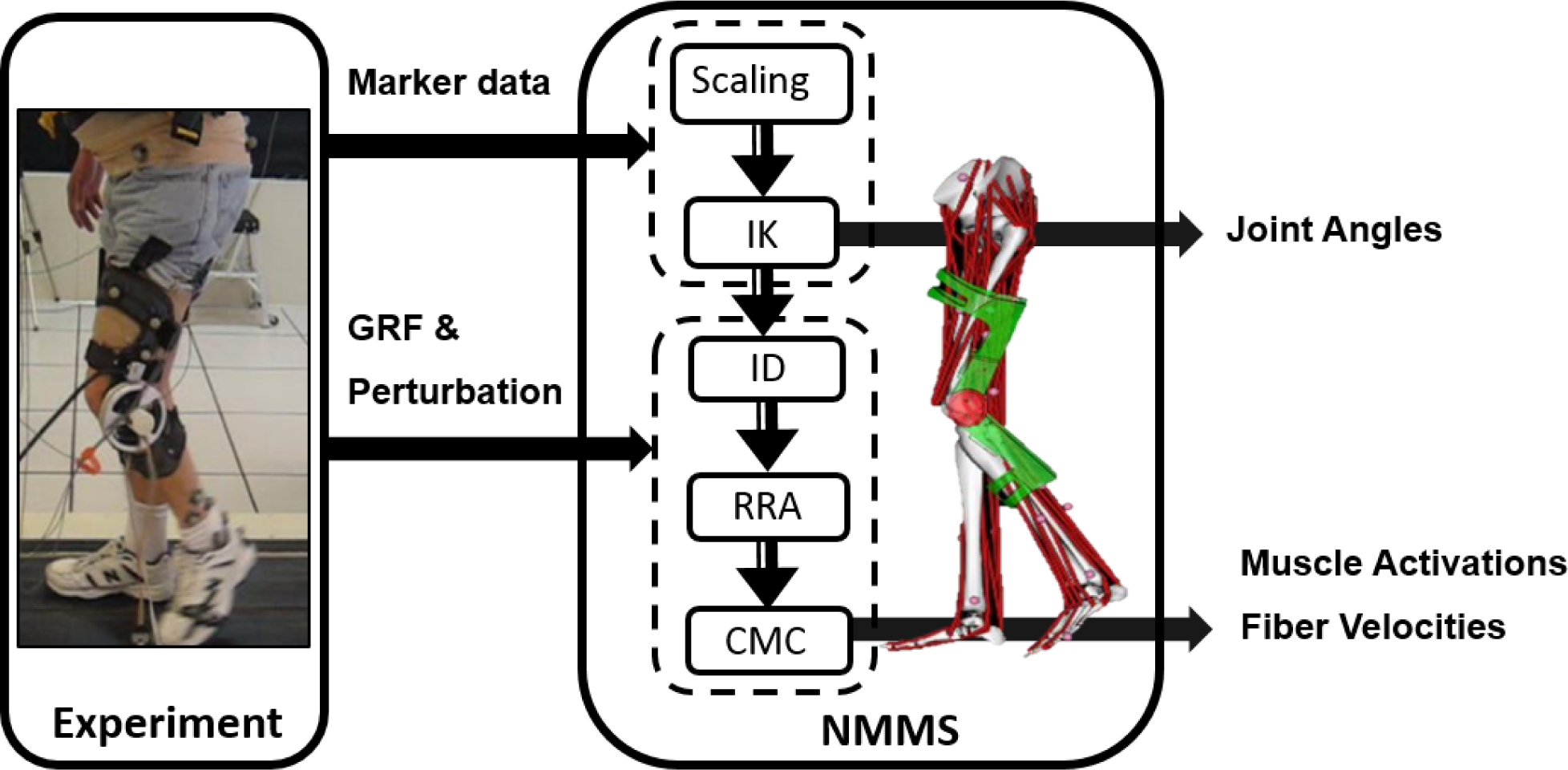
Neuromusculoskeletal modeling and simulation (NMMS) framework. Experimental data is used as input to determine joint angles, and muscle length, velocity and activation profiles. The NMMS processes are shown in the solid frame on right. Height adjusted models are generated for each participant using the scaling tool and joint angles are calculated using inverse kinematics with the least square fit for marker trajectories. The muscle activation and fiber velocity profiles are determined using computed muscle control (CMC), and validated with the experimentally measured EMG data.

The knee flexion perturbation induces opposing torques on the shank and thigh (Sulzer, Roiz et al. 2009), which were modeled as coupled forces on the tibia and femur in OpenSim. The coupled forces were oriented in opposite directions and set equidistant from the center of mass of the corresponding bodies. We validated the accuracy of the perturbation by comparing the simulated torque at the knee with and without the modeled perturbation to the measured torque from the device. In addition to validating knee flexion perturbation values, we also accounted for unmeasured handrail forces using estimation procedures previously described (Akbas and Sulzer 2015). Some adjustments and constraints were made to the model in order to more accurately represent the experimental conditions. For instance, in the experiment, the left thigh markers were attached directly to the device. The compliance of the interface between the brace and the leg reduced the accuracy of the measured knee flexion angle, thus we reduced the weightings of those markers accordingly. The dynamic consistency of the simulations was evaluated by examining the resulting residual forces and moments (Table S2) and verifying them with the OpenSim guidelines (Hicks, Uchida et al. 2015).

We used CMC to generate muscle activations and muscle fiber stretch velocities from the muscle-driven simulation based on the subject’s movement. To validate the simulated muscle activations from CMC, we used the EMG data collected from RF, AL, VL, and semimembranosus (SM). We qualitatively compared the estimated muscle activities and iEMG for both the healthy controls and those with SKG with and without perturbation. We confirmed the activations in the model occurred during the appropriate time in the gait cycle by comparing the simulated muscle activations with the measured iEMG activity on each participant (Figure S1).

Our primary hypothesis (H1) was that the exaggerated abduction in those with SKG is a reflex coupling between the quadriceps and abductors. The reaction to a reflexive stimulus would occur within 120 ms following onset, including mono- and polysynaptic mechanisms (Pierrot-Deseilligny and Burke 2005). This time interval has been implemented for detecting reflexive responses from quadriceps following mechanical perturbation during gait (Mrachacz-Kersting, Lavoie et al. 2004). We defined the stimulus onset, the initiation of the involuntary response, as the timing of the peak stretch velocity. We calculated the stretch velocity of the quadriceps muscle group (RF, VL, VM and vastus intermedius: VI) by taking the time derivatives of the fiber length measures obtained from the simulation. Then we evaluated all the estimated muscle activities during the involuntary response (IR) period within 120 ms after stimulus onset indicated by simulated peak fiber stretch velocity, integrated over a ±1% gait cycle window around the peak. Correspondingly, the voluntary muscle response (VR) was evaluated within the voluntary time window, 120-300 ms following the stimulus onset.

It is also feasible that a previously unaddressed excessive RF activity alone could produce sufficient abduction motion during gait since the muscle has a small abduction moment arm. To test this possibility, we used the musculoskeletal model and modified the RF activation to the maximum value (100%) between the pre-swing and mid-swing phases. We also implemented a partial forward dynamics simulation of a representative stroke subject with the applied perturbation. We forced the simulation to follow the motions calculated from the experimental data for all joints except hip abduction of the affected side, which is driven by the simulated muscle activity patterns. We compared three different cases: 1) simulated abductor and RF activations from perturbed gait, 2) simulated abductor and RF activities from baseline, and 3) simulated abductor activity from baseline and full RF activation (100%).

The alternative hypothesis (H2) was the voluntary knee flexion-hip abduction coupling originated from the lack of independent joint control (synergies), exaggerated as an adaptation to the perturbations. The specific mechanism could be a coupling between hamstrings and abductors (Cruz and Dhaher 2008). In this case, we focused on a different set of steps when the perturbation was temporarily removed (catch trials), expecting a coupling between abductor and hamstring activity dependent on gait adaptation (Reinkensmeyer, Wynne et al. 2002). To test this, we integrated simulated hamstrings activities, including semimembranosus (SM), semitendinosus (ST) and biceps femoris long head (BFlh) within the peak abductor activation regions (±2% gait cycle) for the baseline and catch trials.

### 2.3 Statistical Analysis

The hypothesized reflex coupling (H1) requires a reflex followed by a coupling of the homonymous (i.e., quadriceps) and heteronymous (i.e., abductor) muscle activation (Finley et al., 2008). Quadriceps reflex was determined both experimentally and through simulation. Experimental evidence of quadriceps reflex was provided by quadriceps iEMG within 120 ms following stimulus onset. We used repeated-measures analysis of variance (ANOVA) (*α* < 0.05) with 2 levels: baseline and perturbation, followed by Tukey-Kramer post hoc testing to compare quadriceps iEMG between groups (healthy controls and SKG). We predicted a significant interaction effect between group and condition, with RF iEMG being higher in those with SKG compared to the healthy controls.

Simulation was also used to verify quadriceps reflex. We implemented a linear mixed-effects model, taking groups and fiber stretch velocities as covariates, and dependent variables of simulated quadriceps (RF, VL, VM and VI) activations. We predicted a significant correlation between simulated RF activation in the IR period and its fiber stretch velocity in those with SKG (*α* < 0.05). Lastly, the coupling between quadriceps and abductor activity was investigated using a linear mixed model with peak simulated quadriceps activation as the independent variable and abductor muscles (GMed, GMax, GMin, TFL, sartorius (SAR) and piriformis (PIR)) as dependent variables during IR period. We predicted a significant correlation between RF and abductor activity in SKG compared to the healthy controls.

A voluntary abnormal coordination pattern (H2) would be represented by coupled activation regardless of perturbation. A linear mixed-effects analysis was used with factors of condition (catch trials and baseline) and groups (healthy and SKG). We examined all combinations of simulated hamstrings activity occurring simultaneously with peak simulated abductor muscle activations (GMax, GMin, GMed and TFL) as dependent variables. We expected to find a significant correlation between hamstring and abductor activation in those with SKG compared to the healthy controls. For all linear mixed-effect models subjects were included as random variables. All statistical analyses were performed using R software (R Development Core Team, 2008) using a linear mixed model (lme4) package.

## 3 RESULTS

### 3.1 RF Reflex Activity (H1)

Representative data show increased simulated RF activations and EMG measures in response to the perturbation of an impaired individual, with no such effect on healthy control (Figure 2 A-D). These effects were similar at the group level. We observed a significant interaction effect across groups (healthy controls and SKG) and conditions (baseline and perturbation) in RF iEMG (*F*_*(1,321)*_*=15.5, p<0.001*) but no significant interaction was observed for VL iEMG (*F*_*(1,321)*_*=3.79, p=0.06*). Between knee perturbations and baseline trials, RF iEMG increased significantly (*F*_*(1,235)*_*=86, p<0.001*). The increased RF activity was consistent across subjects (Figure 3A). Similarly, there was a significant decrease in VM iEMG (*F*_*(1,235)*_*=8.01, p=0.005*), but this increase was not consistent across subjects (Figure 3B). For healthy controls, we found a significant decrease in VL iEMG (*F*_*(1,86)*_*=9.04, p=0.003*) whereas no significant changes were observed in VL iEMG (*F*_*(1,86)*_*=0.26, p=0.614,* Figure 3c) or RF iEMG (*F*_*(1,86)*_*=0.02, p=0.89*).

**Figure 2.**
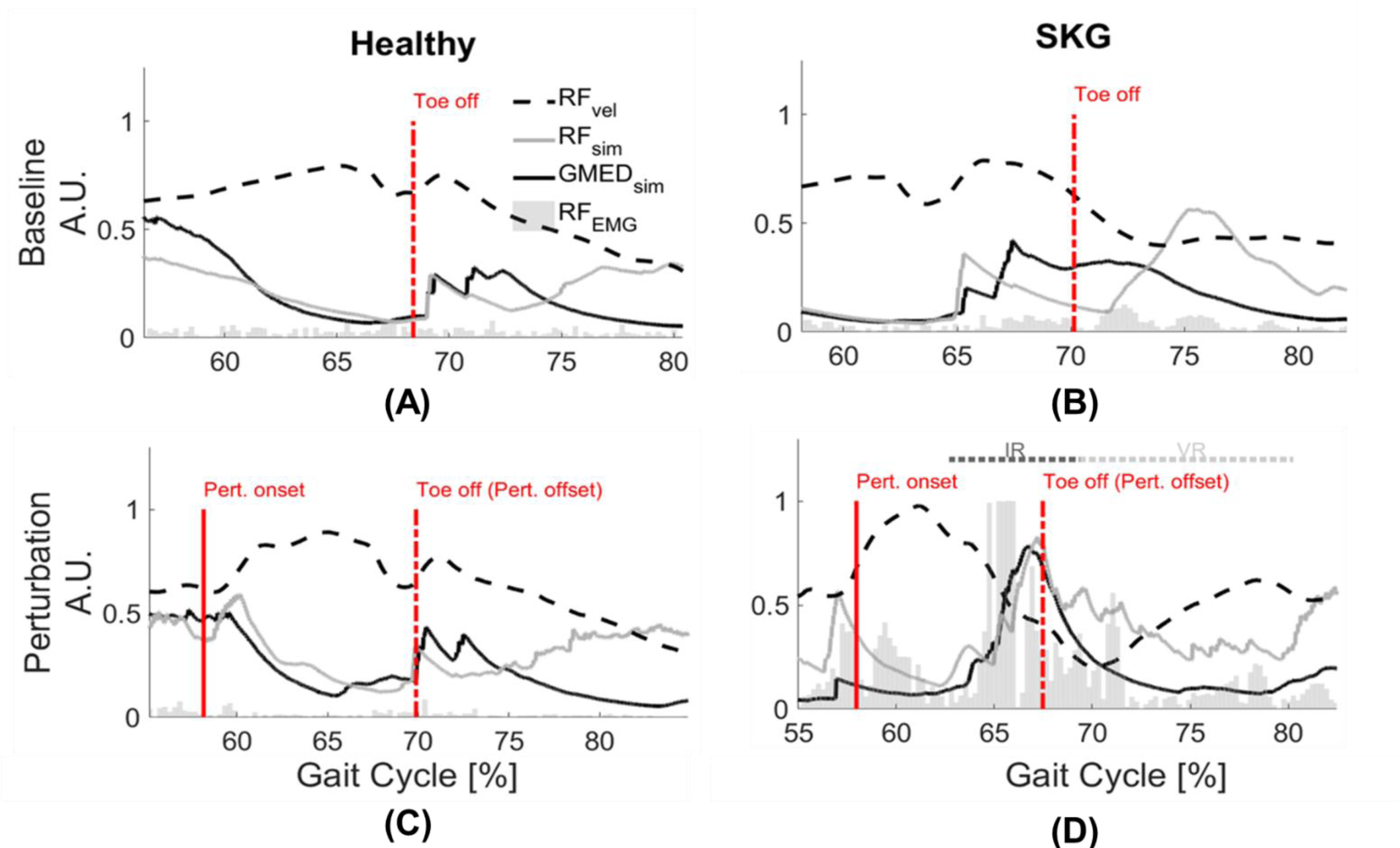
GMed and RF activations are increased with RF stretch velocity in stroke. Rectified RF EMG, simulated RF fiber stretch velocity and muscle activations of RF and GMed for a representative healthy control (left) and individual with SKG(right) with and without perturbation. Each gait interval begins in pre-swing (55-60%gait cycle) and continues until late-swing (80-85% gait cycle). The RF fiber velocities were normalized with respect to the maximum of the demonstrated steps and RF EMG measures are normalized with respect the maximum values throughout the gait cycles of the corresponding subjects. The RF fiber stretch velocity values are increased for both healthy and SKG individuals with the perturbation. However, the raw RF EMG measures and simulated GMed and RF activity only increased in the individual with SKG following peak RF fiber velocity. This is indicative of an abnormal coupling pattern between RF and GMed in post-stroke SKG. The activations of RF and GMed are increased during the involuntary response (IR) period (<120 ms).

**Figure 3.**
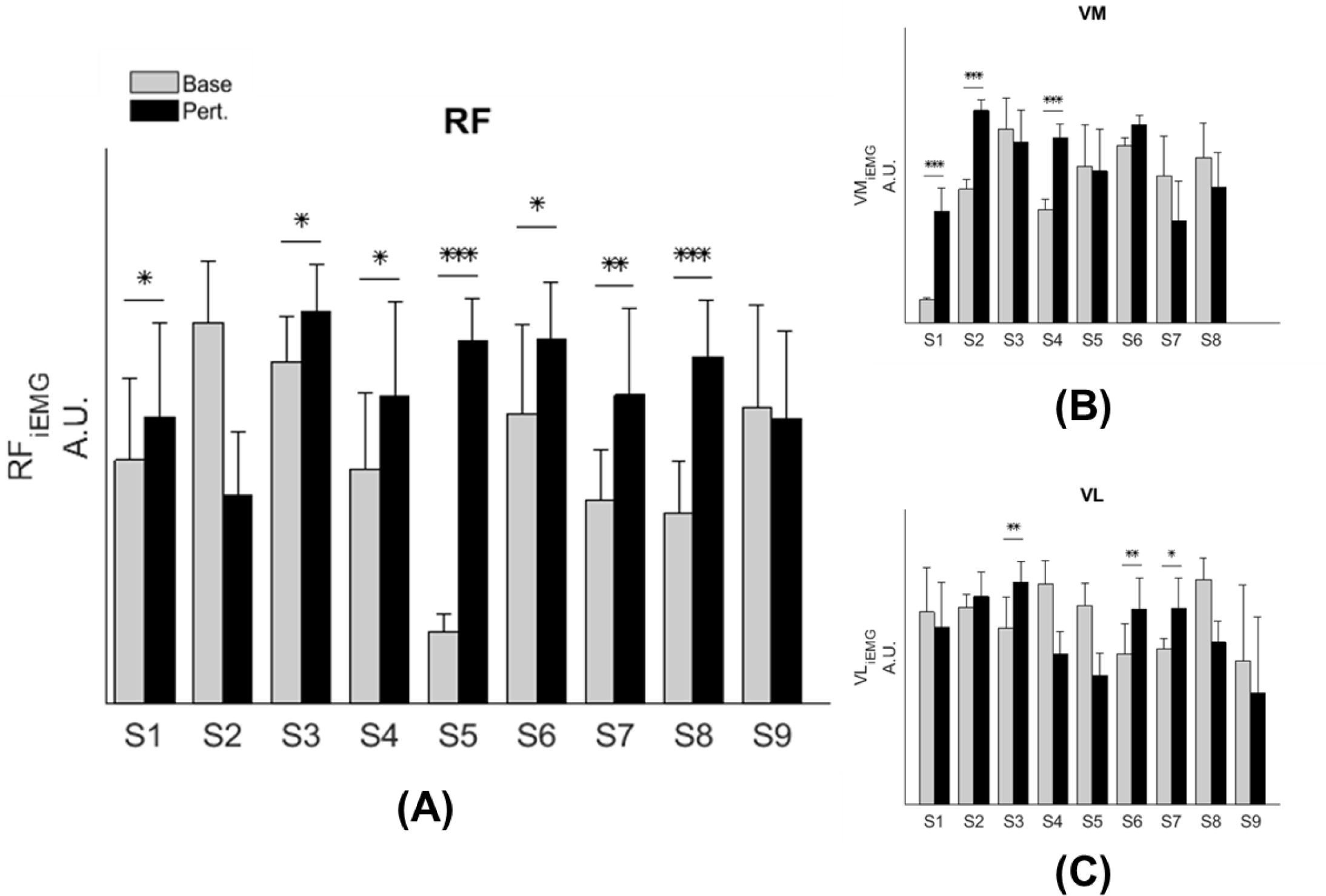
RF activity increases during knee flexion perturbation consistently in people with SKG. RF iEMG (A) and VM iEMG (B) values were increased significantly within the SKG group between baseline and perturbation (p <.001, p<.005 respectively). No significant changes were observed in VL iEMG (C) values (p =.06) based on the repeated measures ANOVA. The increase was more consistent for RF iEMG values across subjects compare to VM iEMG values. “Base” refers to unperturbed Baseline steps, whereas “Pert.” refers to steps with torque perturbations during pre-swing. Significant differences within subjects was determined using Tukey-Kramer post-hoc testing (*p < 0.05, ** p < 0.01, *** p < 0.001).

Simulation results were consistent with EMG measures following the perturbation. Figure S1 shows general agreement between measured and computed muscle activations. We also verified the dynamic consistency of the simulation with the experimental measures kinematic and kinetic and accuracy of the simulated muscle actuation with residual forces and moments from RRA tool and reserved actuators from CMC tool (Table S2) according to the guidelines of OpenSim (Hicks, Seth et al. 2015). Figure 4 shows every modeled step from all subjects and subsequent regression lines derived from the linear mixed model. Figure 4A illustrates peak modeled RF activation following increased RF fiber stretch velocity (RF_vel)_ during the involuntary response period; RF activation increases with RF_vel_ in the SKG group (β_SKG_ = 0.8) but slightly decreases in the healthy controls (β_Healthy_ = −0.03). Compared to healthy controls, people with post-stroke SKG had a significantly greater correlation between simulated RF_vel_ and RF activation (*F*_*(1,321)*_ *=30.2, p=0.001*, Figure 4A). The effect was consistent across SKG subjects (Table S1).

**Figure 4.**
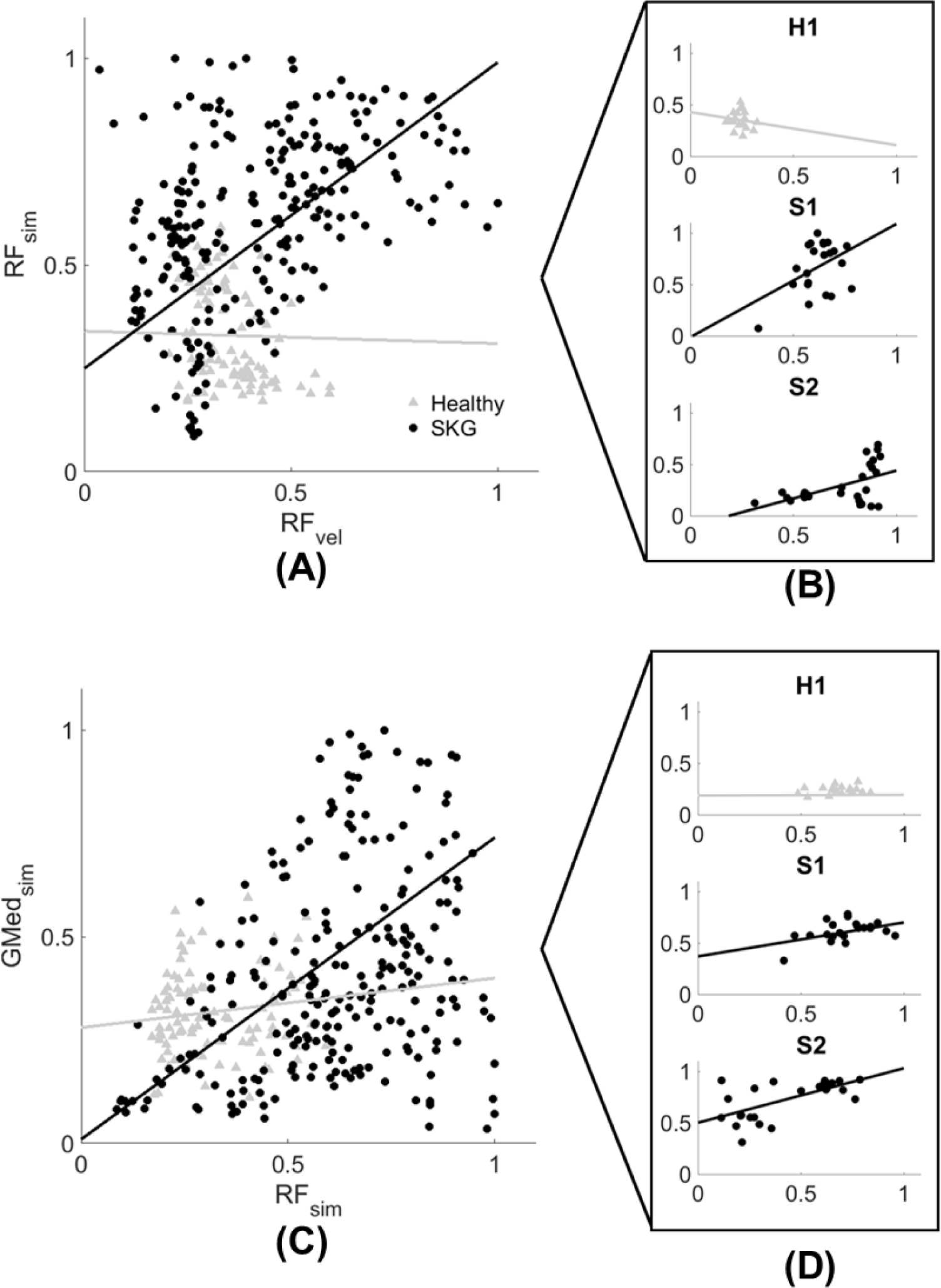
Computed RF activation increases with RF_vel_ (A-B) and coupled with GMed (C-D) activity in people with SKG. Normalized RF peak stretch velocity (RF_vel_) values and peak RF activation (A) and coupled GMed activity (C) from every simulated gait cycle in all subjects during the involuntary response period. Individuals with SKG (dark circles) show increased peak RF (β_SKG_ = 0.8) activity and coupled GMed (β_SKG_ = 0.73) activity, as would be expected in a reflex response. However, healthy controls (gray triangles) show a decrease in RF (β_Healthy_ = −0.03) and small increase in coupled GMed (β_Healthy_ = 0.12) activity, suggesting that this abnormal reflexive response did not exist in healthy controls. A representative healthy participant (H1) and two representative subjects with post-stroke SKG (S1, S2) highlight the correlation between RF_vel_ and RF activation profiles (B) and coupled GMed activity (D).

We found no significant difference between correlations during the voluntary response period (*F*_*(1,321)*_*=0.59, p=0.443)*. We did not find a significant correlation between the activation of other quadriceps muscles during the involuntary response period with increased RF_vel_ (VL: *F*_*(1,321)*_*=0.31, p=0.57*, VM: *F*_*(1,321)*_*=0.66, p=0.412*, VI: *F*_*(1,321)*_*=2.81, p=0.092*). The increased RF activation applied to the generic Gait 2392 musculoskeletal model resulted in less than 2° increase in hip abduction, which was less than the average increase (5°) observed in those with post-stroke SKG. The result of forward simulation with full RF activation in the representative subject with post-stroke SKG resulted in greater hip abduction than baseline (3°), which was 3.5° lower than the abduction following the mechanical perturbation (7.5°).

### 3.2 Reflex Coupling with Abductors (H1)

Representative data show no coupling between simulated RF and GMed activation in the unimpaired individual (Figure 2b). However, when perturbed, the individual with SKG exhibited co-activation of RF and GMed within the involuntary response period. At the group level, simulated GMed activation was correlated to peak RF activation (F(1,321)=7.15, p<0.001, Figure 4C) within the involuntary response period in SKG subjects. This is further illustrated in Figure 4C for all steps and subjects, the correlation between GMed and RF activations were high in those with SKG (β_SKG_= 0.73), but not for the healthy controls (β_Healthy_ = 0.12). The effect was consistent across seven of nine individuals with SKG (Table S1). We found significant correlations of GMax (F(1,321)=5.90, p=0.015) with RF activation in those with SKG, but not with other abductors, GMin (F(1,321)=1.95, p=0.162), TFL (F(1,321)=3.02, p=0.083), SAR (F(1,321)=0.20, p<0.65) or PIR (F(1,321)=0.034, p=0.85) (Table 1). There was no evidence of abductor activity in the voluntary response period following peak RF_vel_ (p>0.05).

**Table 1.** Couplings following perturbation in post-stroke gait. The table shows the level of significance (* p < 0.05, ** p < 0.01) between the corresponding simulated muscle activation profiles and peak RF activation within involuntary response period (*x*_1_) and given the fixed effects of groups (*x*_2_). The results indicate a significant increase between RF and gluteal muscle (GMed and GMax) activation profiles for individuals with SKG compared to the healthy controls during the involuntary response period. No significant difference found for other abductor muscles (GMin, TFL, SAR and PIR) during this period.

| Muscle | $p_{x_1 \times x_2}$ |
| --- | --- |
| GMed | 0.0079** |
| GMax | 0.015* |
| GMin | 0.16 |
| TFL | 0.08 |
| SAR | 0.65 |
| PIR | 0.85 |

### 3.3 Voluntary Synergistic Coupling (H2)

Simultaneous activation of hamstrings and abductors could be part of an abnormal synergistic coupling. We found no significant differences in any combination of simulated hamstrings (SM, ST and BFlh) and corresponding peak abductor (GMed, GMin, GMax, TFL, SAR, PIR) activations compared between conditions (catch trials and baseline) and groups (*p*>0.05).

## 4 DISCUSSION

Previous research found exaggerated hip abduction in individuals with post-stroke SKG with applied knee flexion perturbations (Sulzer, Gordon et al. 2010). We have proposed two possible abnormal neuromuscular mechanisms that could explain the increased hip abduction: a voluntary synergistic coupling between hip abduction and hip extension or an abnormal reflexive coupling between the quadriceps and abductor(s). The measured EMG and NMMS data both indicate that the flexion perturbation produced an abnormal stretch reflex in RF. This activity was coupled simultaneously with simulated GMed activation, but not the other non-gluteal abductors, suggesting an abnormal RF-GMed reflex coupling (H1). These results present a new mechanism of abnormal reflex coordination in post-stroke gait.

These findings also challenge earlier assumptions of hip circumduction being part of a compensatory motion to account for reduced knee flexion (Perry and Burnfield 1992). Our observations suggest that while motions such as vaulting and pelvic obliquity may compensate for lack of knee flexion, circumduction in SKG may be due to other factors such as abnormal coordination patterns. It should be noted that circumduction can be a necessary compensation for other disabilities such as foot drop, but for those with SKG, excessive plantarflexion is either non-existent or alleviated with an ankle-foot orthosis. Regardless of the level of ankle dysfunction in our participants, our data showed abnormal coordination between abductors and RF following knee perturbations. These abnormal patterns may be initiated by excessive RF activity, which is one of the suggested causes of SKG after stroke (Kerrigan, Gronley et al. 1991, Goldberg, Anderson et al. 2004, Lewek, Hornby et al. 2007, Reinbolt, Fox et al. 2008). This initial evidence indicates that interventions intending to restore healthy, symmetric gait for those with SKG should consider hip circumduction as a discoordination pattern rather than a compensation. In addition, clinical interventions that stretch RF, including body-weight supported treadmill training (Hesse, Bertelt et al. 1995) or robotic assistance (Mayr, Kofler et al. 2007) should examine the potential consequences of such an approach.

The simulated RF-GMed coupling appears to be unique. We found no evidence of a stretch reflex in the VL, VI or VM in those with SKG. However, we did find coupled simulated activation of the GMax in addition to GMed. We estimated the individual contributions of abductor muscles during the simulation to address the specificity of the abductor muscles involved in the coupling. For each abductor muscle, we removed the corresponding muscle from the model and allowed a reserved hip abductor/adductor actuator to replace the contribution of the moment generated by the removed muscle. The highest moment (> 40 Nm) was generated by the reserved actuator after the GMed was removed followed by GMax with the second highest contribution (< 10 Nm) indicating the essential role of GMed. These results suggest that GMed is the most likely contributor to excessive hip abduction observed in SKG with knee perturbations.

The neural mechanism underlying this involuntary coupling remains unclear. The malfunction in supraspinal control following stroke is suggested to cause impairment in the regulation of relevant spinal interneurons (Dietz and Berger 1984). This impairment could influence the medium latency stretch responses believed to be relayed through oligosynaptic pathways (Corna, Grasso et al. 1995) which could initiate abnormal heteronymous stretch reflex responses in stroke patients. Previous research has identified the existence of these abnormal reflex responses in static postures. For instance, increased soleus H-reflex excitations have been observed with simultaneous peroneal and femoral nerve stimuli suggesting the excitatory heteronymous pathways in stroke compared to healthy controls (Dyer, Maupas et al. 2009). Others have observed that mechanical perturbations of the AL resulted in a stretch reflex coupling with RF in stroke patients (Finley, Perreault et al. 2008). From our experimental data, we observed those with post-stroke SKG exhibit an increased RF EMG activity consistently within a latency following torque perturbation indicative of a hyperactive stretch reflex. This effect was not observed in EMG measures from VL and VM. The simulated RF activation was closely aligned with measured RF EMG activity as well as simulated simultaneous RF and GMed activation and consistent across subjects. The simulated vasti muscle activations were not aligned with abductor activations following perturbation indicating that the increased RF is initiating the abnormal involuntary response in abductors. We can only speculate whether the RF-GMed coupling is a spinal reflex as the temporal accuracy of simulated muscle activations is not sufficient during gait (> 10 ms), thus further experimental evidence is needed to establish this mechanism.

While NMMS provides rich information about the neuromusculoskeletal system, it has some limitations. Musculoskeletal modeling and simulation rely on a number of assumptions, including cadaver-based muscle moment arms (Menegaldo, de Toledo Fleury et al. 2004), generic Hill-type muscle models and mathematically driven cost functions (Thelen and Anderson 2006). Since our aim was to characterize the neuromuscular mechanism behind the observed response using the simulation, we have not modified the neural control portion of the model with a stretch reflex controller (DeMers, Hicks et al. 2017) or models of spasticity (Van der Krogt, Seth et al. 2013, Jansen, De Groote et al. 2014) to avoid manipulating the muscle states. As a result, simulations require validation with measured muscle activity. An alternative approach could be to use EMG-informed (Demircan, Khatib et al. 2009) or EMG-driven (Rajagopal, Dembia et al. 2016) simulations. Due to the inaccessibility of surface EMG for small abductor muscles and the limited number of total EMG measures in the experiment, we were not able to implement these approaches. However, our simulations were dynamically consistent with experimental force and position measures and were in good agreement with the EMG data of participants (Figure S1). In addition, the reflexive coupling between RF and GMed were consistent for the participants with post-stroke SKG (Figure 4, indicated by S1 and S2) with the highest and lowest correlation for simulated activations and RF EMG (Figure S1, indicated by S1 and S2). Furthermore, both simulated and experimental RF results showed consistent effects within and between subjects. Thus, our data suggest that simulated muscle activities are indeed realistic.

## 5 CONCLUSION

Based on previous work (Sulzer, Gordon et al. 2010), our simulation analysis suggests that knee flexion perturbations in pre-swing initiate a hypersensitive reflex response in RF in those with post-stroke SKG. The stretch reflex of RF coupled with simultaneously activated GMed suggests a previously unidentified heteronymous abnormal reflex coordination during gait. This represents new evidence of abnormal reflex coordination during gait in people with post-stroke SKG. Our results suggest that hip circumduction may not be entirely a compensatory motion, but rather part of an abnormal coordination pattern. Such information could influence how therapists treat hip circumduction post-stroke, from treating compensations to treating coordination. However, further work is needed to more accurately determine the nature of this abnormal coordination pattern in stroke and its effects on function.

