## Supplementary figures and tables for "Neuromusculoskeletal simulation reveals abnormal rectus femoris-gluteus medius reflex coupling in post-stroke gait"

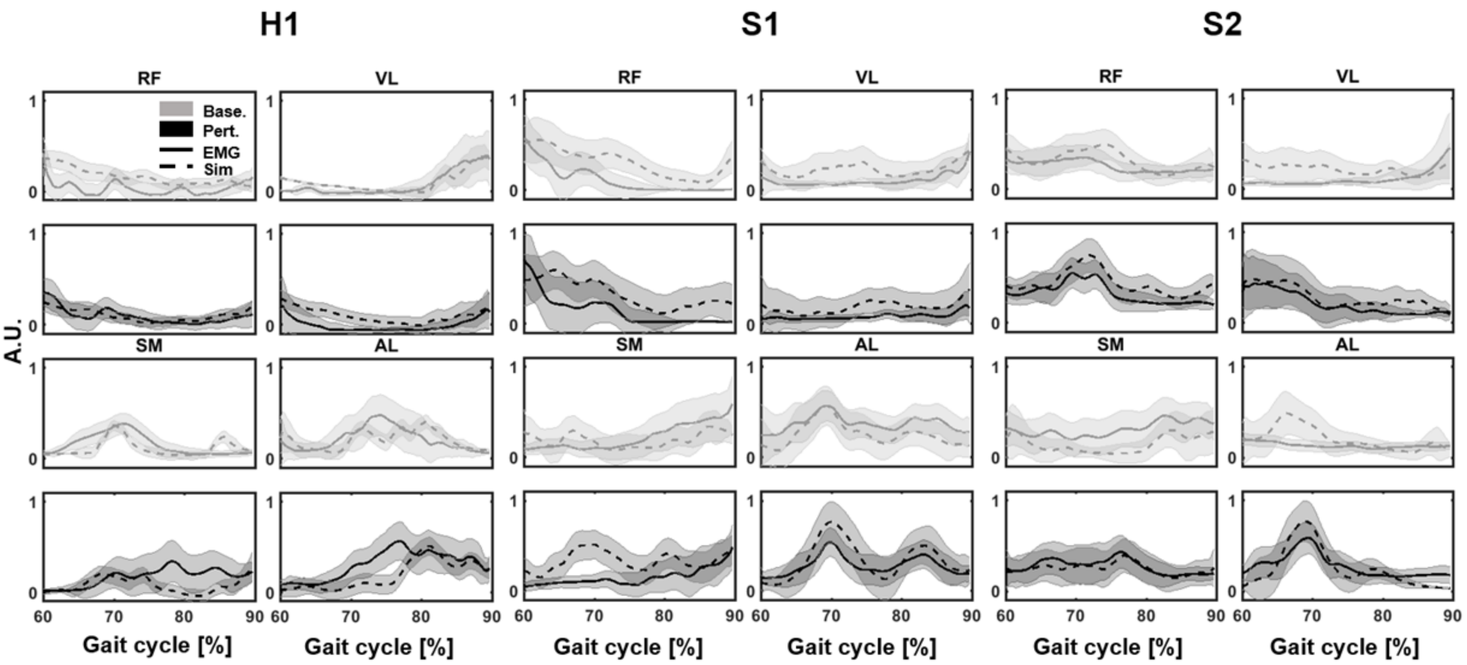


**Figure S1: Validation of the simulated muscle activities (dashed lines) with integrated EMG activity (solid lines) for baseline (gray) and perturbation (black) for a representative healthy participant (H1) and two representative participants with post-stroke SKG (S1 and S2).** The figure represents qualitative comparison between the simulated muscle activations and integrated EMG values for a healthy participant and two stroke participants with lowest and highest total Pearson correlation (baseline: $r_{S1}$ = .42, $r_{S2}$ = .72; perturbation: $r_{S1}$ = .46, $r_{S2}$ = .76), sum of the correlation values for baseline and perturbation, between RF EMG and simulated RF measures respectively during pre-swing and late-swing. EMG measures are collected from RF, vastus lateralis (VL), adductor longus (AL) and semimembranosus (SM). The healthy participant is in good agreement both for the baseline and perturbation conditions but there is a larger variance between participants and steps with post-stroke SKG variance observed by the different profiles and variance within participants indicated by shaded areas (± SE).

**Table S1. Correlations between $RF_{vel}$ and simulated muscle activations of RF and between RF and GMed activations in people with SKG during the involuntary response period.** The table shows the correlation coefficients and significance (* p < 0.05, **p < 0.01, *** p < 0.001) between $RF_{vel}$ and simulated RF (left) and, RF and GMed activations (right) during IR for each participant with SKG.

| Subject | $r_{RF_{vel}*RF}$ | $r_{RF*GMed}$ |
|---|---|---|
| S1 | $0.41^{***}$ | $0.31^{**}$ |
| S2 | $0.58^{*}$ | $0.95^{**}$ |
| S3 | $0.95^{***}$ | $0.48^{**}$ |
| S4 | $0.23^{***}$ | $1.35^{***}$ |
| S5 | $0.38^{***}$ | $0.75^{***}$ |
| S6 | $0.56^{***}$ | $0.72^{***}$ |
| S7 | $0.33^{**}$ | 0.1 |
| S8 | $-0.2$ | 0.38 |
| S9 | $1.08^{***}$ | $1.61^{***}$ |

**Table S2.** Maximum and average values of the residual forces ($F_x$, $F_y$, $F_z$) and moments ($M_x$, $M_y$, $M_z$) from reduced residual algorithm (RRA) tool and the corresponding reserved joint actuators with maximum torque values from computed muscle control (CMC) tool of OpenSim are shown in the table. The values are collected across all simulated for each participant from the region of the gait cycles used in data analysis. **The maximum values are within the range of good simulation based on OpenSim guidelines (J. Hicks, Seth, Hamner, & Demers, 2015) for each participant with post-stroke SKG.**

| | **Residual Forces (N)** | | | | | | **Residual Moments (N*m/s)** | | | | | | **Reserved Actuators (N*m/s)** | |
|---|---|---|---|---|---|---|---|---|---|---|---|---|---|---|
| | $F_x$ | | $F_y$ | | $F_z$ | | $M_x$ | | $M_y$ | | $M_z$ | | Corresponding reserved joint actuator | Max. |
| | Avg. | Max. | Avg. | Max. | Avg. | Max. | Avg. | Max. | Avg. | Max. | Avg. | Max. | | |
| S1 | -.62 | 14.4 | .35 | 8.23 | 1.7 | 11.4 | -.14 | 26.5 | -1.8 | 11.6 | -2.4 | 28.5 | Left ankle flexion/extension | 22.9 |
| S2 | 1.9 | 8.77 | 1.6 | 8.93 | 1.3 | 6.54 | .38 | 11.5 | 2.6 | 10.7 | 1.6 | 26.9 | Left ankle flexion/extension | 19.2 |
| S3 | 1.4 | 9.65 | 1.8 | 5.07 | 1.2 | 12.7 | 0.2 | 13.5 | -1.1 | 12.9 | -.05 | 20.0 | Right knee flexion/extension | 17.0 |
| S4 | .84 | 6.38 | -.48 | 7.62 | -1.2 | 5.18 | -.44 | 17.7 | -3.8 | 9.61 | .02 | 35.7 | Left ankle flexion/extension | 16.9 |
| S5 | 2.3 | 14.4 | .14 | 13.0 | -.21 | 10.8 | -.58 | 18.7 | -1.7 | 17.0 | 1.5 | 28.5 | Left ankle flexion/extension | 24.1 |
| S6 | 2.1 | 6.06 | 1.4 | 14.1 | -.61 | 6.70 | -.19 | 24.4 | -3.3 | 18.3 | -1.1 | 26.0 | Right knee flexion/extension | 20.8 |
| S7 | 2.2 | 12.4 | -.33 | 6.74 | -1.6 | 9.29 | -.02 | 25.3 | .72 | 19.5 | .13 | 33.9 | Left hip flexion/extension | 22.7 |
| S8 | 2.0 | 10.6 | .25 | 6.33 | .73 | 7.30 | -1.1 | 29.1 | -4.2 | 15.5 | -2.8 | 30.9 | Left ankle flexion/extension | 26.6 |
| S9 | 1.4 | 5.67 | 1.8 | 2.50 | 1.4 | 4.45 | .20 | 27.9 | -.14 | 18.7 | -.22 | 25.0 | Right knee flexion/extension | 22.2 |
